# Reasoning by exclusion in food-caching Eurasian jays (Garrulus glandarius)

**DOI:** 10.1101/2024.10.14.618234

**Authors:** Isobelle Hawkins, Michael Huemer, Susan Carey, Nicola S. Clayton

## Abstract

Two related methods have been widely used to test animals’ capacity to reason by exclusion: Call’s (2004) 2-cups task, where subjects choose between two cups in which one reward was hidden and the empty cup is revealed; and Premack & Premack’s task (1994) where one reward goes in each of the two cups and one reward is removed. In both cases, success is identifying the cup with the reward. It has been suggested that among corvids, the demands associated with caching may foster exclusion abilities by reinforcing the experience of relevant contraries (e.g., empty vs. full), or by supporting domain general capacities that support caching and exclusion inferences. However, Shaw et al. (2013) found that Eurasian jays, proficient cachers, failed a version of Call’s 2-cups task. To further test the caching hypothesis, we amended the procedure using invisible displacement to hide rewards, removing aspects of Shaw et al.’s procedure that may have masked competence of Eurasian jays. We tested seven Eurasian jays, who showed spontaneous success on both Call’s 1-reward/show-empty (80% correct) and Premack and Premack’s 2-reward/remove-1 (85% correct). These results align with the hypothesis that corvids’ exclusion reasoning is related to caching behaviour.

## Introduction

Reasoning by exclusion has been extensively studied as it may provide a window into the logical capacities of non-human animals (Tomasello & Call, 1997). The most common method deployed is Call’s (2004) 2-cups task. In this task, animals encounter two opaque cups, a reward hidden in one of them. Without knowing which one is baited; the animal is shown that one cup is empty. A related task, from Premack and Premack (1994), is to hide two equally desirable rewards, one in each of the two cups, then remove one reward. In both tasks, success is selecting the baited container. Both tasks involve excluding one of the cups as a possible location of a reward.

Völter & Call (2017) reviewed data from these tasks across 33 species. Spontaneous success (i.e., from the first trial without training) was exhibited by great apes, some other primate and corvids also achieved spontaneous success but at a slightly lower rate than great apes. Furthermore, there is evidence that suggests the rates of spontaneous success among corvids may be ordinally related to their caching behaviour (see Figure 1). For instance, non-caching jackdaws perform at chance on the 2-cups task (Schloegl et al., 2011). In contrast, Clark’s nut- crackers, skilled cachers, relying upon it for survival, are capable of storing up to 33,000 seeds annually (Vander Wall & Balda, 1977), and excel at this task, outperforming all other birds tested (Tornick & Gibson, 2013; Völter & Call, 2017). Ravens, though less prolific cachers (de Kort & Clayton, 2006; see Table 1, pp. 418), also succeed, though not as consistently as nut- crackers (Schloegl et al., 2009; Tornick & Gibson, 2013). Carrion crows, who cache less intensely than these other corvids and do not rely on caching for survival (Goodwin, 1976), did not show spontaneous success and passed only after multiple trials and after controlling for local enhancement cues (Mikolasch et al., 2011).

**Figure 1.**
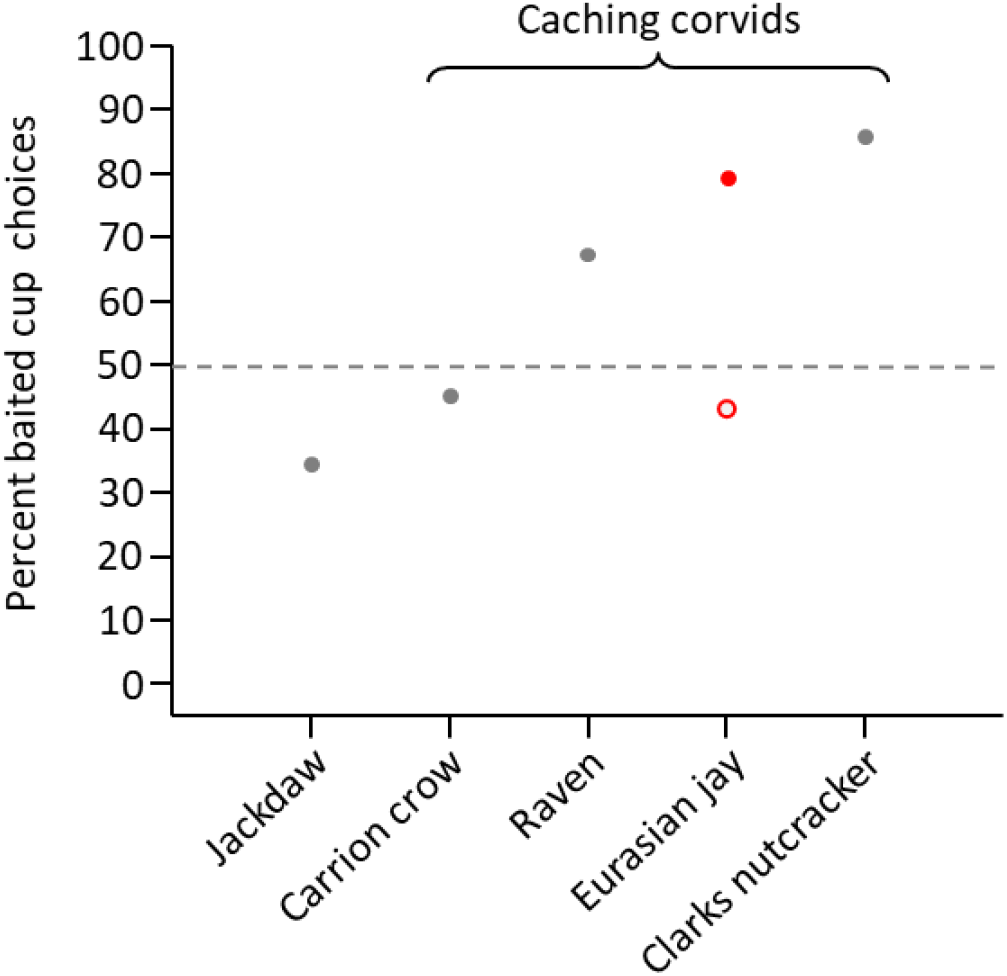
Mean percent of choosing the baited cup in the 1-reward/show-empty task. On the x-axis, the corvids are arranged with the least prolific cachers on the left and the most prolific cachers on the right (Tornick & Gibson, 2013; Vander Wall, 1990). The open red circle indicates Eurasian jays’ performance from Shaw et al. (2013), the red dot shows performance in the current study. Dashed line indicates 50% chance level.

Given this ordinal relationship, Mikolasch et al. (2011) and Schloegl et al. (2009) hypothesised that the good performance of some corvids derives from their food-caching behaviour. The cognitive demands of caching, such as tracking the location of caches and avoiding sites already recovered or pilfered, may enhance exclusion abilities as it promotes the experiences of relevant contraries (i.e. “empty” vs. “full.”). Contraries are a form of logical negation; it cannot be that both contraries are true of a single state of affairs (see Bermudez, 2003, 2006). Challenging this hypothesis are data from Eurasian jays depicted on Figure 1, open red circle. Eurasian jays are intense and proficient cachers, relying upon it for survival (Clayton et al., 1996), falling between ravens and Clark’s nutcrackers with approximately 11,000 acorns caches a year (Vander Wall, 1990), yet the only study of a 2-cups task with this species found failure (38% correct, Shaw et al. 2013). This failure is surprising given that Eurasian jays have shown other advanced general cognitive abilities such as tool selection tasks that involve predictive or forward inference (Cheke et al., 2011; Davidson et al., 2017), which would make them candidates for success in the 2-cups task. It is therefore possible that the unexpected failure of Eurasian jays in this task is a false negative.

Shaw et al. (2013) used a modified procedure of Call’s 2-cups task. Unlike in other versions of the task, the experimenters turned around and baited the cups out of sight. The Eurasian jays might have been unsure whether the reward was actually deposited into one of the cups, and thus did not engage in reasoning they were otherwise capable of. This procedure involved significant movement both when moving the cups in and out of the testing compartment and when one of the cardboard lids was removed to reveal the empty cup. The present study tests whether changing these procedural details reveals spontaneous success on both the 1-reward/show-empty and the 2-reward/remove-1 tasks. We explore whether the level of success falls between that of ravens and Clark’s nutcrackers. We changed the procedure in two ways (see Figure 2). *Firstly*, we familiarised the Eurasian jays to the cups and the movements during training trials (which gave no training in reasoning by exclusion). *Secondly*, we did not use novel objects e.g. occluders or lids as Eurasian jays are known to be neophobic. Instead, we used an invisible displacement procedure to hide the rewards giving unambiguous evidence that worms were deposited into cups. Engelmann et al. (2022) found that chimpanzees’ performance in 4-cups tasks with occluders and with invisible displacement did not differ.

**Figure 2.**
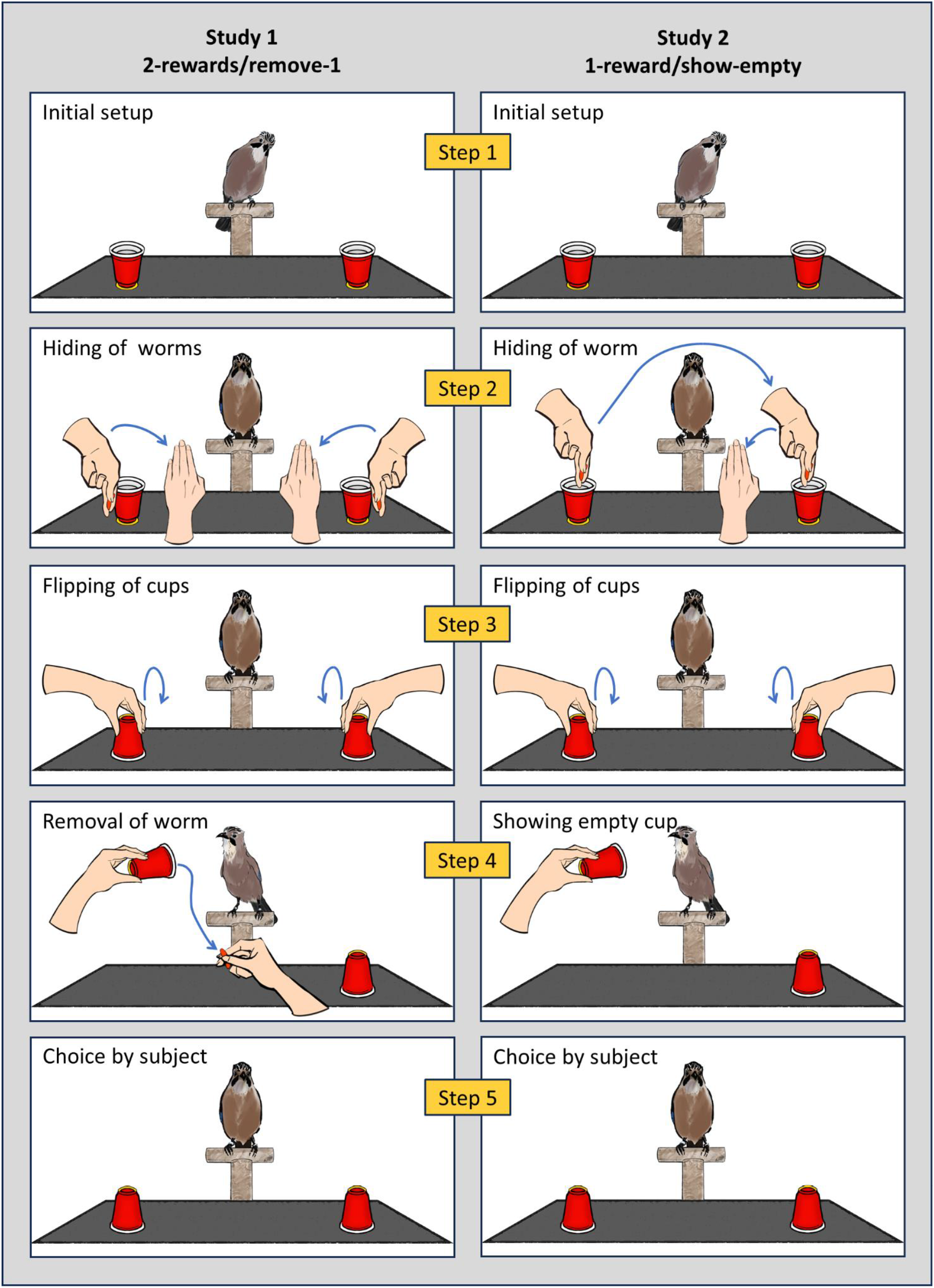
Procedure of Studies 1 and 2. Worms are depicted in red for visibility.

No study we know of directly compares the two versions of the 2-cup exclusion task within the same participants. We do so here to explore whether both tasks tap into a common exclusion capacity.

**Study 1: *2-rewards/remove-1***

## Methods

### Subjects

Seven Eurasian jays (four females; aged 9 years) participated. One additional subject was tested (Booster, male) but only participated for two of the 20 test trials and was excluded from data analysis. The subjects were hand raised and lived in a group of 14 Eurasian jays, in an aviary, where they extensively cached. See Supplement for details of the housing and participation.

### Design & procedure

Subjects were tested individually. Before testing, each subject was habituated to the experimental setup by familiarising them to picking up and revealing the contents of a single cup. The Eurasian jays received four training trials and two blocks of ten test trials; all subjects but Booster completed 20 test trials. See Supplement for a description of the materials, the training trials, the counterbalancing and randomisation of training and test trials.

### Test trials

The Eurasian jays were shown that the cups were empty (Figure 2, left side, Step 1). The rewards were then placed on opposite sides of the A3 piece of black cardboard (Step 2), the cups were then flipped on top of the rewards (Step 3). Then in full view one reward was removed from a cup (Step 4). The subject was then allowed to pick one cup (Step 5).

## Results and discussion

The Eurasian jays chose the baited cup 85% of the time (see Figure 3A, left side), significantly above chance (one-sample Wilcoxon signed-rank test: *Z* = 2.29, *p* = .022) with a very strong effect size (*r* = .90). This finding suggests that the negative findings from a previous study (Shaw et al., 2013) underestimated the jays’ capabilities for reasoning by exclusion.

**Figure 3.**
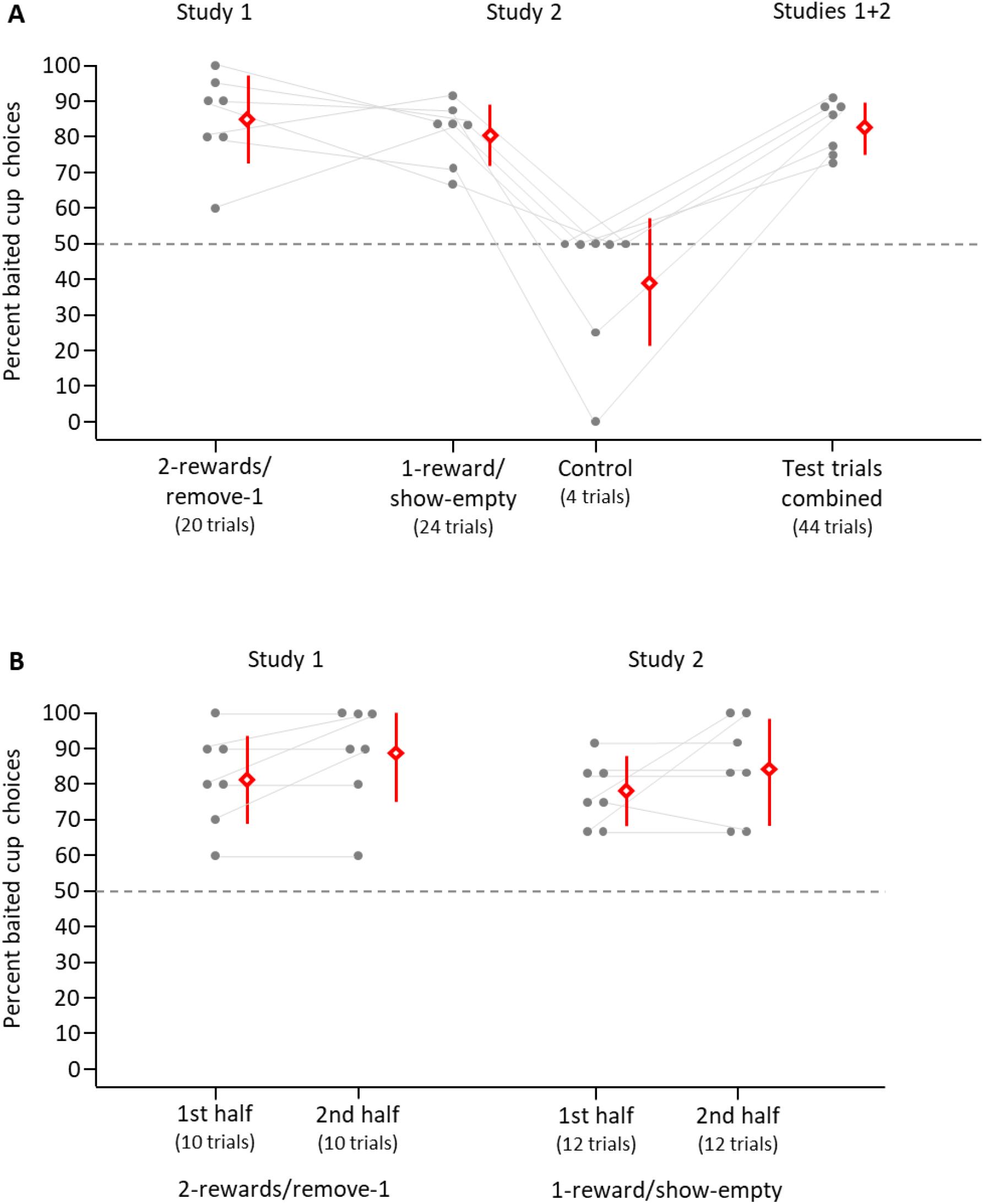
Percentage choice of the baited cup in Study 1 and Study 2. (A) Performance in Study 1 and Study 2. (B) Comparison of performance of the first and second half in Studies 1 and 2. Grey dots indicate an individual Eurasian jay’s percentage of baited cup choices. Red diamonds show means, error bars indicate 95% confidence intervals. Dashed lines indicate 50% chance level.

The success of the Eurasian jays on this task was spontaneous. Without having been exposed to tasks that require reasoning by exclusion, or choosing between two cups to receive a reward, they performed well from the beginning. Performance was good on both the first and second half (ten trials each) of the study (see Figure 3B, left side): On the first ten trials they chose the baited cup 81% of the time (above chance, one-sample Wilcoxon signed-rank test: *Z* = 2.29 *p* = .022, effect size: *r* = .90) and 89% on the second ten trials (above chance, one-sample Wilcoxon signed-rank test: *Z* = 2.39 2.3, *p* = .021, effect size: *r* = .90. There was no significant difference between the first and second ten trials, related-samples Wilcoxon signed-rank test: *Z* = 1.37, *p* = .17. These results rule out the possibility that the Eurasian jays’ performance could have been associative learning over trials.

Eurasian jay’s excellent performance can also be observed on the individual level. In Figure 3 (left side), the grey dots show each subject’s individual performance. One of the Eurasian jays picked the baited cup 60% of the time (not different from chance, binomial test: *p* = .503), while the other six Eurasian jays’ performance was at least 80% (all above chance, binomial test: all *p* ≤ .012). The excellent performance of nearly all the Eurasian jays (six out of seven) on this task demonstrates a robust ability to reason by exclusion in this species.

**Study 2: *1-reward/show-empty***

## Methods

### Subjects and Materials

Study 2 included the same subjects and materials as in Study 1.

### Design and procedure

The Eurasian jays received four training trials and four blocks of six trials per day. All subjects completed 24 test trials. This varied to Study 1 as Eurasian jays looked to cache after around 6 trials. After the test trials were completed, the subjects received one block of four control trials. See Supplement for a detailed description of the training trials, and the counterbalancing and randomisation of training and test trials.

### Test trials

The Eurasian jays first saw that the cups were empty (Figure 2, right side, Step 1). The experimenter hid the reward from view in their fingers then put these fingers into one cup and then the other cup placing the reward into one of the cups (Step 2). After flipping the cups (Step 3), the experimenter showed the Eurasian jay the contents of the empty cup (Step 4). The subject was given a choice of one of the two cups (Step 5).

### Control trials

To ensure that the procedure was not giving away the rewards location (e.g., the Eurasian jays might actually see where the worm is hidden), we presented the Eurasian jays with four control trials. These trials followed the same Steps 1 to 3 (Figure 1, right side) as in the test trials but provided no information about the cup’s contents (i.e., omitted Step 4; Figure 1). The subject was then allowed to choose a cup.

## Results and discussion

Similar to the *2-rewards/remove-1* task in Study 1, Eurasian jays performed well on the *1-reward/show-empty* test trials in Study 2. They picked the baited cup 80% of the time (see Figure 3A, right side), significantly above chance (one-sample Wilcoxon signed-rank test: *Z* = 2.30, *p* =.021), the effect size (*r* = .90) is very strong.

The Eurasian jays picked the baited cup 39% of the time on the *control trials* which is not different from chance, one-sample Wilcoxon signed-rank test: *Z* = 0.89, *p* = .371. Five Eurasian jays chose the baited cup half the time (two of four), and four of these always picked either the right or the left cup, showing a side bias. Having a side bias in the control trials (but not the test trials) confirms that performance is different when the Eurasian jays got no information which of the two cups contained the reward. The Eurasian jays’ good performance in the test trials depends upon the information that is input to an exclusion inference.

Eurasian jays showed spontaneous success on the test trials of Study 2, just as in Study 1. Performance was high on both the first and the second half (twelve trials each) of the study (see Figure 3B, right side): Eurasian jays picked the baited cup on the first twelve trials 78 % of the time (above chance, one-sample Wilcoxon signed-rank test: *Z* = 2.29, *p* = .022, effect size: *r* = .90) and on the second twelve trials 85% of the time (above chance, one-sample Wilcoxon signed-rank test: *Z* = 2.29, *p* = .022, effect size: *r* =.90). There was no significant difference between the first and second twelve trials, related-samples Wilcoxon signed-rank test: *Z* = 1.07, *p* = .500. Again, this speaks against Eurasian jays’ success being based on associative learning over trials.

Eurasian jay’s individual performance on this task was again robust. In Figure 3A (middle panel), the grey dots show each Eurasian jay’s individual performance. Five of the seven Eurasian jays picked the baited cup 83% of the time or more (all above chance, binomial test: all *p* ≤ .002). One further Eurasian jay was at 71% (marginally significant above chance, binomial test: *p* = .064), and the remaining Eurasian jay was at 67% (not different from chance, binomial test: *p* = .152).

When combining Studies 1 and 2 (see Figure 2A, right side), Eurasian jays performed well. Picking the baited cup 83% of the time, significantly above chance (one-sample Wilcoxon signed-rank test: *Z* = 2.29, *p* = .022), the effect size (*r* = .90) is very strong. On the individual level, each Eurasian jay picked the baited cup at least 73% of the time (i.e., all seven were all above chance by the binomial test: all *p* ≤ .004). The Eurasian jay who picked the baited cup at 60% in Study 1 performed excellently in Study 2 (83%) and the Eurasian jays who were at 67% and 71% in Study 2 performed excellently in Study 1 (90% and 80%, respectively).

In sum, the results from Study 2 converge with those of Study 1 in demonstrating that Eurasian jays are capable of spontaneous reasoning by exclusion. In Study 1, Eurasian jays’ success demonstrated that (1) they can update their beliefs when shown that a reward is removed from its location, and, thus, the location was then empty, and (2) eliminate that location as a place for searching for that reward. In Study 2, the Eurasian jays had to eliminate a possible location and restrict their search for the reward to the other possible location (the second cup) after they were shown the contents of the empty cup. A final analysis showed performance on the two different tasks did not significantly differ (related-samples Wilcoxon signed-rank test: *Z* =1.35 *p* =0.18), consistent with the conclusion that each tapped into a common capacity for exclusion inference.

### General discussion

Collectively, Eurasian jays chose the target cup 80% or more (significantly above chance) both when shown the removal of a reward from a cup when each cup had been baited (Study 1) and when shown that one of the cups was empty when only one cup had been baited (Study 2). The Eurasian jay’s performance was spontaneous, showing a strong preference for choosing the target cup from the beginning, and every individual’s performance was better than chance when the data from both studies were combined. Eurasian jays spontaneously reasoned by exclusion. However, it is important to highlight a key difference between the two studies. In Study 2, there is ambiguity about the location of the rewards after the hiding. Showing a cup to be empty allows an inference that resolves that ambiguity. In contrast, there is no ambiguity about the location of the rewards in Study 1; Study 1 did not require an inference about an unknown location. Nonetheless, both tasks involved ruling out one of the cups as a possible location of a reward. The similarity in results indicates that this shared element is driving the observed variance in the data (i.e., variance around 80% correct in each procedure).

The Eurasian jays’ performance stands in contrast to previous results found by Shaw and colleagues (2013) who found no evidence that Eurasian jays could reason by exclusion. We conclude that procedural features of their implementation masked the Eurasian jay’s capacity for spontaneous exclusion inferences. When these procedural features were changed in the present study, the Eurasian jays’ level of spontaneous success fell between that of ravens and that of Clark’s nutcrackers (see Figure 1, red dot), as predicted by the hypothesis that spontaneous reasoning by exclusion is predicted by a species caching proficiency. Further research should systematically explore this relationship. The procedures and sample sizes in the studies reported on Figure 1 were slightly different. This relationship could be established empirically, by testing different species on the same tasks, where individuals have had identical previous experience with choice tasks.

Further research should adjudicate between two broad, not mutually exclusive, explanations for the relation between caching proficiency and success at exclusion inferences. *First*, there may be evolutionary support for domain general cognitive capacities that support exclusion inferences and supporting caching. That is, cognitive demands associated with caching (e.g., working memory, spatial memory, episodic-like memory, and future planning) may also give rise to the capacity for exclusion (Mikolasch et al., 2011; Schloegl et al., 2009). *Second*, extensive caching experience may provide opportunities for learning necessary for the exclusion tasks. One proposal is that non-human animals may not possess a domain-general operator for “not” that they can draw upon in exclusion inferences (Bermudez, 2003). Instead, they may base their exclusion reasoning on specific contraries like “empty” and “contains food”. That specific contraries may need to be learned, pair by pair, might explain why many adults of species tested require many training trials (sometimes hundreds), before passing the 2-cups task (Bohn et al., 2020). Caching behaviour may provide input sufficient for learning this specific contrary. For instance, cachers frequently encounter pilfering and empty cache sites. An empty food location may inform a cacher about the fate of previously present food, a scenario non-caching species do not face (Grodzinski & Clayton, 2010). Alternatively, caching behaviour may exercise domain-general capacities relevant to exclusion, making animals with extensive caching experience more likely to draw upon those capacities spontaneously in the 2-cups tasks. The hypothesis that caching behaviour supports relevant learning could be tested by controlled rearing studies, testing whether extensive caching behaviour is needed for exclusion inferences. For example, one could test whether individual crows, rooks or ravens, who have never cached either spontaneously succeed on the 2-cups tasks, or learn faster than do individuals from species that never cache, as a function of the place on ordinal scale of their species.

Some researchers have interpreted success on the 1-reward/show-empty task as evidence for the capacity for reasoning through a deductive syllogism (A or B, not A, therefore B) in non-human animals (e.g., referring to it as “causal-logical inference” Call 2004). This requires setting up premises with the content the food is in the right cup or the left cup; it’s not in the left cup, and concluding, it is in the right cup. We agree that the 2-cup success reflects eliminating a search option, and indeed, by 17 to 19 months of age, human children show a capacity for eliminating an option when shown one of two containers in the search space is empty. The question though, is how the search space is being represented. Further research, using a 3-cups or a 4-cups procedure (Mody & Carey, 2016), show catastrophic failures by chimpanzees and young children to represent the relation “or” in these tasks (Engelmann, 2022; Grigoroglou et al., 2019; Leahy et al., 2022; in revision; Mody & Carey, 2016). We illustrate with the 3-cups procedure. A prize is hidden in a singleton cup, and a prize is also hidden in a pair of cups. The participant is given only one chance to get a prize. If they can represent “a prize is in a singleton; a prize is in the left or in the right of the pair, they should reliably choose the singleton cup. Instead, chimpanzees and older 2-year-olds choose one of the pair 50% of the time; and 3-year-olds 40% of time (Engelmann, 2022; Grigoroglou et al., 2019; Leahy et al., 2022; in revision; Mody & Carey, 2016). Leahy et al. (2022) establish that this failure in children is a failure to represent “The prize is in A or B” which is logically equivalent in this situation to “There is one prize in these two cups. The prize might be in A and it might be in B.”. Without one of these premises, the inference the child or the primate is making in the 2-cups task does not have the structure of a disjunctive syllogism. A high priority for future research is to test corvids on the 3- and 4-cups tasks.

## Declarations

### Ethics approval

Experiments were previously approved as a non-regulated procedure by the University of Cambridge’s Animal Welfare and Ethical Review Body, Sub-Standing Committee (Ethics codes: NR2023/52) and followed Home Office Regulations and the ASAB’s Guidelines for the Treatment of Animals in Behavioural Research and Teaching.

## Acknowledgement

We would like to thank Maggie Dinsdale, Jamie Clayton, Georgia Davies, Laura Jenkins, and Harvey Booth, the technicians responsible for the birds’ husbandry, as well as James Davies for his assistance during the early stages of bird training in the project. We would also like to thank Peiru Chen for her efforts in creating the intricate drawings of birds, cups and hands we used to illustrate the procedures of Studies 1 and 2 in Figure 2.

## Data availability

Raw data and all data generated or analysed during this study are included in this published article and its supplementary files.

## Credit author statement

**IH:** Conceptualisation, methodology, formal analysis, investigation, data curation, writing - original draft, writing - review & editing, visualisation. **MH:** Conceptualisation, methodology, formal analysis, data curation, writing - original draft, writing - review & editing, visualisation, supervision. **SC:** Conceptualisation, writing - original draft, writing - review & editing. **NSC:** Conceptualisation, resources, writing - original draft, writing - review & editing, supervision, funding acquisition.

