## Supplement - methods for "Reasoning by exclusion in food-caching Eurasian jays (Garrulus glandarius)"

**Studies 1 and 2**

**Subjects**

Seven adult Eurasian jays were tested from one aviary (four females: Jaylo, Stuka, Soijka and Penny; three males: Homer, Godot, and Poe, aged 9 years old). When not being tested the Eurasian jays lived in one social group in a large outdoor aviary each measuring 20×6×3m (length x width x height). Aviaries were wood and mesh constructions with a gravel floor. Smaller indoor testing compartments (3×1×2m) were connected to the aviary, accessible via opaque hatch doors (0.5×0.5m) and were used for testing; subjects participated voluntarily. The Eurasian jays were hand-raised by members of Clayton's comparative cognition lab especially Dr Rachael Miller, having been obtained from a registered breeder (Mark Ghobbian, Derbyshire, UK) under a Natural England Licence to Professor Nicola Clayton and kept for future experiments after the conclusion of the study.

On test days, the subjects' maintenance diet (a mix of boiled eggs, vegetables, fruit, and seeds with soaked cat biscuits) was removed from the aviary one hour before testing to enhance motivation for coming to the testing room. The Eurasian jays were restricted for a maximum of four hours.

These Eurasian jays had undergone extensive cognitive testing, and several had previously participated in tool use experiments (e.g. Amodio et al., 2019) as well as perception experiments (e.g. Garcia-Pelegrin et al., 2021; Schnell et al., 2021). They had also had exposure to some of the apparatus used in the current experiment e.g. the cups used as hiding locations were used in a previous study (e.g. Schnell et al., 2021). These Eurasian jays had not previously been tested on any inference tasks using either auditory or visual information, with the animals having little or no experience in specifically two-way choice tasks or auditory choice-based tasks.

All experiments were approved by the University of Cambridge (Ethics codes: NR2023/52) and followed Home Office Regulations and the ASAB's Guidelines for the Treatment of Animals in Behavioural Research and Teaching.

### Materials

The experiment was conducted on a wooden board which was placed in the indoor aviary 0.7m above the ground. Two identical red plastic cups were used as hiding locations, (4.5 cm in diameter and 5 cm in height) and a wooden perch (20 cm in width, 14 cm in height) free to move on the wooden board was used to present the set-up (see Figure 2 for initial set up). A string was connected to the bottom of the cups to create a hook so that the Eurasian jays could use their bill to pick the cups up and reveal their contents. The cups were displayed on a black piece of card (29.7 cm x 42 cm), which served as a contrast to the rewards (wax-worms). The cups were presented on opposite sides of the card (32cm apart). Rewards were waxworms, a highly desirable goal for the jays which they were highly motivated to receive. Trials were recorded on a GoPro Hero4 video camera, a small number of files were lost while transferring from the memory card to the hard drive, but all data was written down.

### Study 1: ***2-rewards/remove-1***

### Procedure and design

The trials in Study1 took place between October 2023 to January 2024. Each Eurasian jay was tested individually once a day for around 20 minutes. They received one block of ten trials per day, across two days. If a Eurasian jay lost motivation and ceased approaching the cups, occasionally a block of trials was split across two sessions within a single day. All the Eurasian jays completed 20 test trials.

The Eurasian jays were isolated in a single testing compartment so that they were physically and visually removed from the other Eurasian jays in the aviary. The trials began when the Eurasian jay sat on the perch facing towards the experimenter for at least five seconds, showing no indication of restlessness. Before testing, each subject was individually habituated to the experimental setup by familiarising them to picking up and revealing the contents of a single cup to avoid neophobic reactions during trials. Training trials followed the sequence LRRL. The test trials in study 1 and 2 were all randomised in which cup the reward went into.

### Training trials

Prior to the test trials the Eurasian jays undertook four training trials to ensure that they could reliably keep track of a baited cup. Each trial began with the experimenter showing the subject that the two cups were empty. The cups were placed upside down, and the subjects were shown the reward, then, in full view of the subject, a worm was placed under one cup. Subjects were then allowed to choose one of the cups by picking it up and revealing what was underneath. Only subjects who chose the baited cup three out of the four times moved on to the test phase; all Eurasian jays met this criterion.

### Study 2: ***1-reward/show-empty***

### Procedure and design

The trials in Study 2 took place between December 2023 to January 2024. The Eurasian jays received four blocks of six trials per day. This change was made because we noticed in Experiment 1 that after around six trials the Eurasian jays looked to cache and became distracted. If a Eurasian jay lost motivation and ceased approaching the cups, occasionally a block of trials was split across two sessions within a single day. All Eurasian jays completed 24 test trials. After the test trials were completed, the Eurasian jays received one block of four control trials. As each Eurasian jay had already been familiarised with the equipment from the previous experiment they went straight into training trails.

### Training trials

These training trials ensured that the Eurasian jays could keep track of the baited cup after the reward had been hidden. Each Eurasian jay received four training trials, these trials began by showing the Eurasian jay that the cups were empty, then placing the cups upright on to the black card. The experimenter then showed the Eurasian jay a reward in their palm, then hid the reward in their hand between their thumb, index, and middle finger. The experimenter then put these fingers which held the reward into one of the cups, in full view of the jay. The cup that the experimenter put their fingers into was ordered LRRL across trials. The experimenter then showed the Eurasian jay their palm was empty and then turned the cups upside down so the jay could use the string to pick the cup up. Finally, the experimenter put both their hands over the two cups to avoid stimulus enhancement. After this presentation, the Eurasian jay was allowed to make a choice and received the reward if correct or saw the empty cup if not. Only subjects who chose the baited cup three out of the four trials moved to the test trials, all Eurasian jays met this criterion.
